# Analytical Method Development for Evaluation of Pharmacokinetics and Tissue Distribution of Thiourea-Based Antivirals through nLC/MS-MS

**DOI:** 10.1101/2023.06.28.546831

**Authors:** Jitendra Kumar, Purnima Tyagi, Manisha Yadav, Deepti Sharma, Jaswinder Singh Maras, Vijay Kumar

## Abstract

Thiourea-based antivirals are organosulfur chemical compounds. Due to their various medicinal uses, including antiviral, antioxidant, and anticancer properties, they are becoming more popular. In this study, pharmacokinetics, Metabolism, bioavailability, and distribution of thiourea derivatives to organs of rats. Thiourea derivatives, namely DSA-00, DSA-02, and DSA-09, exhibit characteristics similar to those of conventional drugs when evaluated in terms of pharmacokinetics, drug-likeness, and medicinal chemistry through *in*-silico analysis. Then a simple and sensitive analytical method was developed for quantified DSA-00, DSA-02, and DSA-09, using prednisolone as an internal standard (IS) and nano liquid chromatography tandem mass spectrometry (nLC-MS/MS). The plasma and tissue samples were preprocessing with acetonitrile before chromatographic separation by C18 column with isocratic elution using a mobile phase of water-methanol (30:70, v/v) at a flow rate of 0.6 mL/min. A triplequadrupole tandem mass spectrometer in multiple-reaction monitoring (MRM) scanning via an electrospray ionization (ESI) source functioning in negative and positive mode detection. For DSA-00, DSA-02, DSA-09 and, internal standard the optimized mass transition ion-pairs (m/z) for quantification were 297.2, 311.2, 309.09, and 361.19 respectively. Each analytic run required only 15 minutes between injections. The calibration curve for thiourea derivatives showed good linear regression coefficient (R^2^ >.99) over a range of 1.00-10000 pg/mL except DSA-09. The lower limit of quantification (LLOQ) was 1.00 ng/mL. The intra- and inter-day precisions were no more than 10.8%, and relative errors (RE) ranged from 0.5 to 5.98%. The validated method was effectively used to explore the pharmacokinetics of orally administrative thiourea derivatives in rats.

## 1. Introduction

Thiourea derivatives are a class of organosulfur chemical compounds that have garnered significant interest due to their diverse pharmacological properties (Ronchetti et al., 2021). These properties include anticancer, anti-bacterial, antiviral, anti-fungal, anti-diabetic, anti-malarial, antinociceptive, and other medicinal activities (Abbas, Al-Harbi, and Sh El-Sharief 2020; Ravichandran et al., 2019; Ghosh et al., 2017; Fan et al., 2011; Ullah et al., 2023; Mishra and Batra 2013; Tang et al., 2007). Recent research has further supported the antiviral and antimicrobial properties of thiourea derivatives (Singh et al., 2023; Thanh et al., 2023). These compounds also serve as valuable precursors in the synthesis of acyclic and heterocyclic compounds, facilitating the design of biologically active agents with strong therapeutic potential (Ghorab et al., 2017; Hashem et al., 2020). Moreover, incorporating halogen functional groups into thiourea derivatives has been shown to enhance their biopotency, bioavailability, and lipophilicity (Nunes et al., 2021; Wilcken et al., 2013; Xu et al., 2014). Notably, thiourea derivatives have been utilized in the treatment of various microbiological illnesses, including Mycobacterium tuberculosis, where compounds like p-acetamidobenzaldehyde thiosemicarbazone have been employed for several decades (Ukrainets et al., 2009). Additionally, modifications involving oxygen-containing side chains have demonstrated increased potency against Mycobacterium tuberculosis (Ronchetti et al., 2021).

In a previous study, we introduced a novel antiviral agent, DSA-00 (IR-415), which exhibited promising activity against HBV infection (Ghosh et al., 2017). To enhance its antiviral efficacy, we designed and synthesized twenty derivatives of DSA-00 by making modifications to the parent molecule, such as altering the position of fluorine on the phenyl ring and increasing the carbon chain length between the phenyl thiourea and imidazole moieties (Singh et al., 2023; Kumar et al., 2023 communication). Remarkably, the antiviral efficacy of DSA-00 and its derivatives, namely DSA-02 and DSA-09, was comparable to that of established antivirals like Tenofovir and Entecavir. In cell culture models, two of the newly synthesized derivatives, DSA-02 and DSA-09, exhibited superior effectiveness compared to the parent molecule (Singh et al., 2023; Kumar et al., 2023 communication). Molecular modeling studies validated these findings (Kumar et al., 2023 communication).

The discovery and development of new drugs play a pivotal role in advancing medical research and improving patient outcomes. In recent years, the nLC-MS/MS technique has emerged as a powerful tool in drug discovery, offering unparalleled selectivity, sensitivity, and rapidity (Li et al., 2021). This advancement in mass spectrometry technology has revolutionized the field, enabling precise evaluation of drug metabolism and pharmacokinetics (DMPK) (Ramanathan, 2013; Lai et al., 2022). The nLC/MS technique, in particular, has become the preferred analytical method for DMPK studies due to its efficient coupling with nano liquid chromatography through electrospray ionization (Deschamps et al., 2023). Although numerous studies have evaluated the pharmacokinetics of various compounds using LC-MS/MS after oral administration, there is a lack of research specifically focusing on the pharmacokinetics of thiourea derivatives in vivo.

To address this research gap, we have developed a highly sensitive and rapid nLC-MS/MS method capable of detecting thiourea derivatives in rat plasma following oral administration. Our method not only establishes a new standard for pharmacokinetic evaluation of these compounds but also underscores the potential of the nLC-MS/MS technique in advancing drug discovery and development.

## 2. Materials and Methods

### 2.1. Reagents and chemicals

DSA-00, DSA-02, and DSA-09 were synthesized in our laboratory (purity 98.00%, determined by HPLC). DSA-00, DSA-02, and DSA-09 were fully characterized by ^1^H, ^13^C NMR, and MS data (Singh et al., 2023). As an internal standard (IS), prednisolone (purity 98.99%, determined by HPLC were purchased from Sigma-Aldrich. Merck (Darmstadt, Germany) provided HPLC-grade methanol and acetonitrile. A Milli-Q system (Millipore, MA, USA) was used to prepare deionized water.

### 2.2. *In-silico* identification of Pharmacokinetics, ADME, and Drug likeness

SwissADME (www.swissadme.ch), a web-based tool was used to assess the Pharmacokinetics, drug likeness, and medicinal chemistry behaviors of the thiourea derivatives. According to the Antonie Diana(Daina et al., 2017)

### 2.3. Preparation of standard and quality control (QC) samples

Accurately prepared the standard stock solution of 10 mg/mL DSAs required dissolving the standard in acetonitrile in the proper quantities. Detail’s methods in supporting information.

### 2.4. Method validation, Calibration standards, and Sample preparation

The methods performance was validated in accordance with the FDA 2001 guidelines.(Ponnuru et al., 2012) Calibration standards, QC, and pharmacokinetics sample were used 100 μL plasma samples combined with 100 μL of acetonitrile and 100 □L of IS working solution. The mixture was vortexed for 1 minute before being centrifuged at 10,000 rpm for 5 minutes. The supernatant was then transferred to autosampler vials for injection into the nLC-MS/MS system in the amount of 20μL.

### 2.5. Chromatographic and mass spectrometric conditions

In the chromatographic separation, a C18 column was used, and the mobile phase consisted of a mixture of 30 volumes of solvent A (water) and 70 volumes of solvent B (methanol). The separation was performed using isocratic elution. Detail’s methods in supporting information.

### 2.6. Experimental animal, collection of blood samples, and Pharmacokinetic by MS/MS

Approval was obtained from the Institutional Animal Ethical Committee (IAEC/ILBS/20/09) prior to the study. Six male rats weighing 300 ± 50 g and female rats weighing 200 ± 20 g were obtained from the Department of Comparative Medicine, Institute of Liver and Biliary Sciences, New Delhi, India. Detail’s methods in supporting information.

## 3. Results

### 3.1. *In-silico* Evaluation of pharmacokinetics drug-likeness, physicochemical, medicinal chemistry properties of Thiourea derivatives

In this study, the swissADME online tool was utilized to predict the pharmacokinetics, bioavailability, drug-likeness, and medicinal chemistry friendliness of Thiourea derivatives. The BOILED Egg, iLOGP, and bioavailability radar algorithms were employed to predict ADME parameters, bioavailability, and medicinal chemistry of these derivatives. The results of the study showed that DSA-02 and DSA-09 had higher gastrointestinal absorption rates compared to DSA-00. Additionally, all three derivatives successfully crossed the blood-brain barrier. Furthermore, the study investigated the skin permeation and observed that DSA-02 had a higher negative value (−6.66cm/s) compared to DSA-00 (−6.28cm/s) and DSA-09 (−6.28cm/s). The thiourea derivatives did not exhibit p-glycoprotein substrate activity. The study also investigated the inhibitory effect of these derivatives on cytochrome P450 and found that they inhibited the activity of CYP1A2, CYP2C19, and CYP2D6, except CYP2C9 and CYP3A4. The study predicted the bioavailability of the thiourea derivatives and observed that they all had the same bioavailability score of 0.55. The water-solubility of DSA-00 was less than DSA-02 and DSA-09. In addition, the study calculated five parameters related to the solubility of thiourea derivatives such as Log P o/w (iLOGP), XLOG3P, WLOGP, MLOGP, and SILICOS-IT. Moreover, the drug likeness of thiourea derivatives was investigated using Lipinski rule, Ghose, Veber, Egan, and Muegge filter. All three derivatives obeyed these rules with zero violation. The study also investigated the medicinal chemistry friendliness of these derivatives and found one violation for DSA-02 and DSA-09, while DSA-00 had zero violations. The study also found zero PAINS structural alerts and one Brenk structural alert for all three derivatives. Finally, the study investigated the bioavailability radar for oral bioavailability prediction, which showed a range of >-0.7 and +5 for LIPO as XLOGP3. The SIZE as molecular weight and POLAR as TPSA were also determined (Figure 1). The study found that these thiourea derivatives fell within the pink colored portion of the radar, indicating promising candidates for leading drugs. The study concluded that these thiourea derivatives showed GI absorption and blood-brain barrier penetration, making them potential drug candidates. Furthermore, they were determined to be P-gp-substrate negative, which suggests that they would not interfere with glycoprotein activity (Table S1). *In-silico* pharmacological study reveals that these thiourea derivatives have drug properties, on the basis of these result, performed *in vivo* pharmacological study on male and female rats.

**Figure 1:**
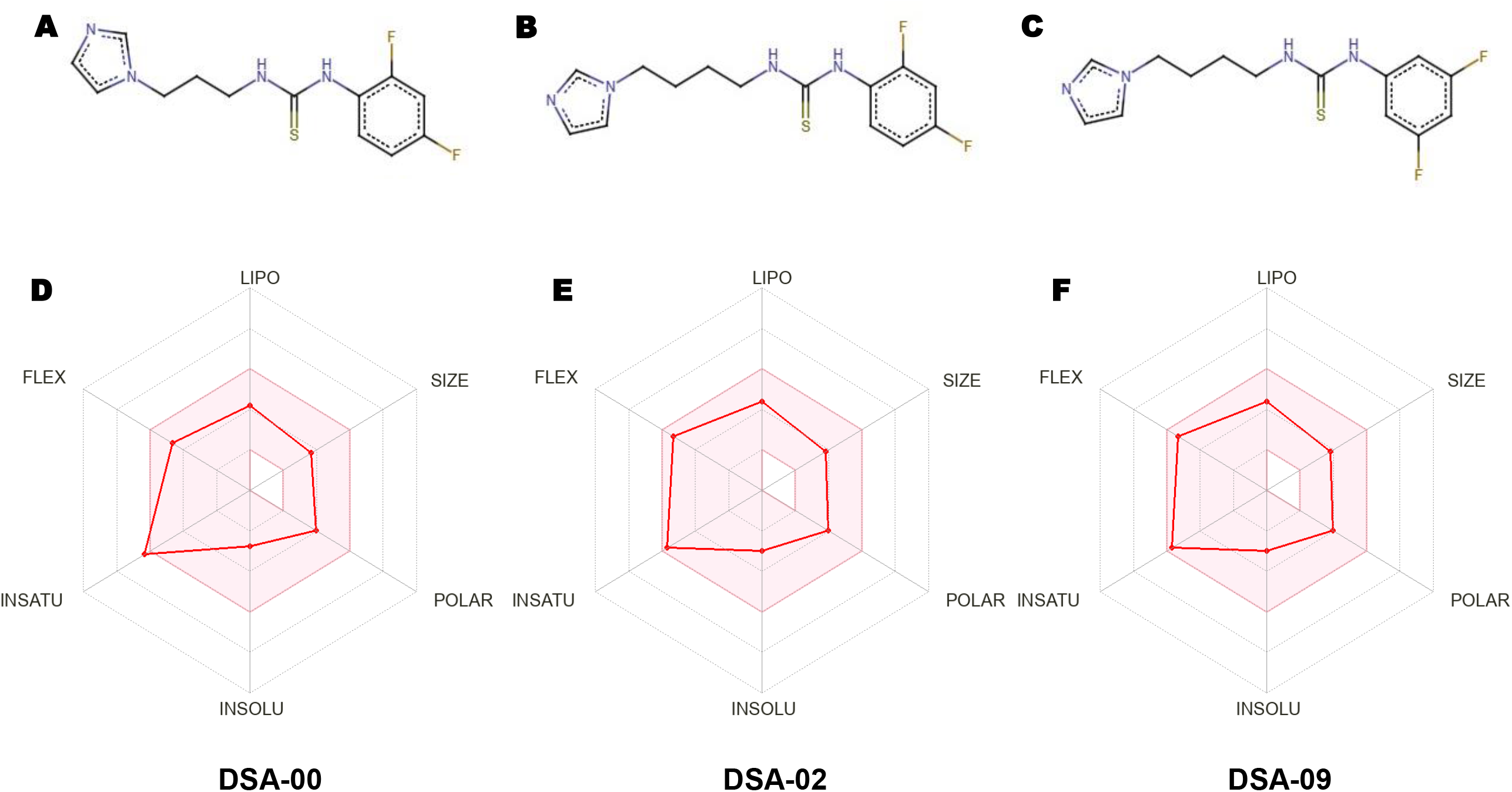
Sketch of chemical structure and Bioavailability radar of thiourea derivatives. The chemical structures of thiourea derivatives (A= DSA-00, B= DSA-02, and C = DSA-09). Bioavailability radar (pink region) for the demonstration of thiourea derivatives (LIPO = lipophilicity as XLOGP3; SIZE = size as molecular weight; POLAR = polarity as TPSA (topological polar surface area); INSOLU = insolubility in water by log S scale; INSATU = insaturation as per fraction of carbons in the sp3 hybridization; and FLEX = flexibility as per rotatable bonds.

### 3.2. Chromatographic separation, Mass spectrometric detection, and Method standardization

This study optimized experimental conditions by directly injecting thiourea derivatives and the internal standard (IS) prednisolone into the mass spectrometer at optimum concentration. Positive ion modes had a superior signal-to-noise ratio than negative ions, except DSA-09. Thiourea derivatives and prednisolone have primary precursor molecular ions at m/z 297.2, 311.2, 309.09 and 361.19, respectively. The product ion scan showed thiourea derivatives most stable product ions at m/z 2311.2, 309.09 and prednisolone at 361.19. Thus, we selected mass transitions of m/z 309.2 -293.4 for DSA-00, 320.3-303-.3 for DSA-02, 320.3-303-.3 for DSA-09 and 387.1 - 328.3 for prednisolone in MRM mode (Figure 2). Therefore, evaluation of specificity and selectively of thiourea derivatives. First tried methanol-water and acetonitrile-water mobile phases to establish optimal chromatographic conditions. Formic acid in the aqueous phase improved sensitivity, while ammonium acetate improved peak shape. Methanol outperformed acetonitrile in shape of peak and background noise. We used mobile phases (A) of 90% methanol and 0.1% formic acid and (B) of water with 5 mmol of ammonium acetate and 0.1% formic acid (90:10, v/v). Thiourea derivatives and prednisolone had the best peak form, less background noise, more intensity, and faster run times. The analytical method specificity and selectivity were assessed for both the internal standard (IS) and thiourea derivatives. For pooled blank rat plasma and liver homogenate samples as well as samples spiked with thiourea derivatives at the Lower Limit of Quantification (LLOQ) level, representative Multiple Reaction Monitoring (MRM) chromatograms were produced. Additionally, chromatograms from tissue homogenate and plasma samples taken an hour after dosing were examined. These chromatograms are shown in Figure 2 and Figure S1-12. It is substantial to notice that neither the internal standard nor thiourea derivatives retention time were observed to be affected by any endogenous metabolites. Moreover, precision and accuracy in the plasma and tissue samples calibration curves for homogenates, in particular, showed strong linearity in the concentration ranges of 10-10000 pg/mL and 1-5000 pg/mL respectively. In rat plasma and tissue homogenates, the Lower Limit of Quantification (LLOQ) for thiourea derivatives was found to be 10 pg/mL and 1 pg/mL, respectively. For performing pharmacokinetic analyses and evaluating the distribution of thiourea derivatives in tissues, these LLOQ values were deemed sufficient. Please see Table S2 for more information.

**Figure 2:**
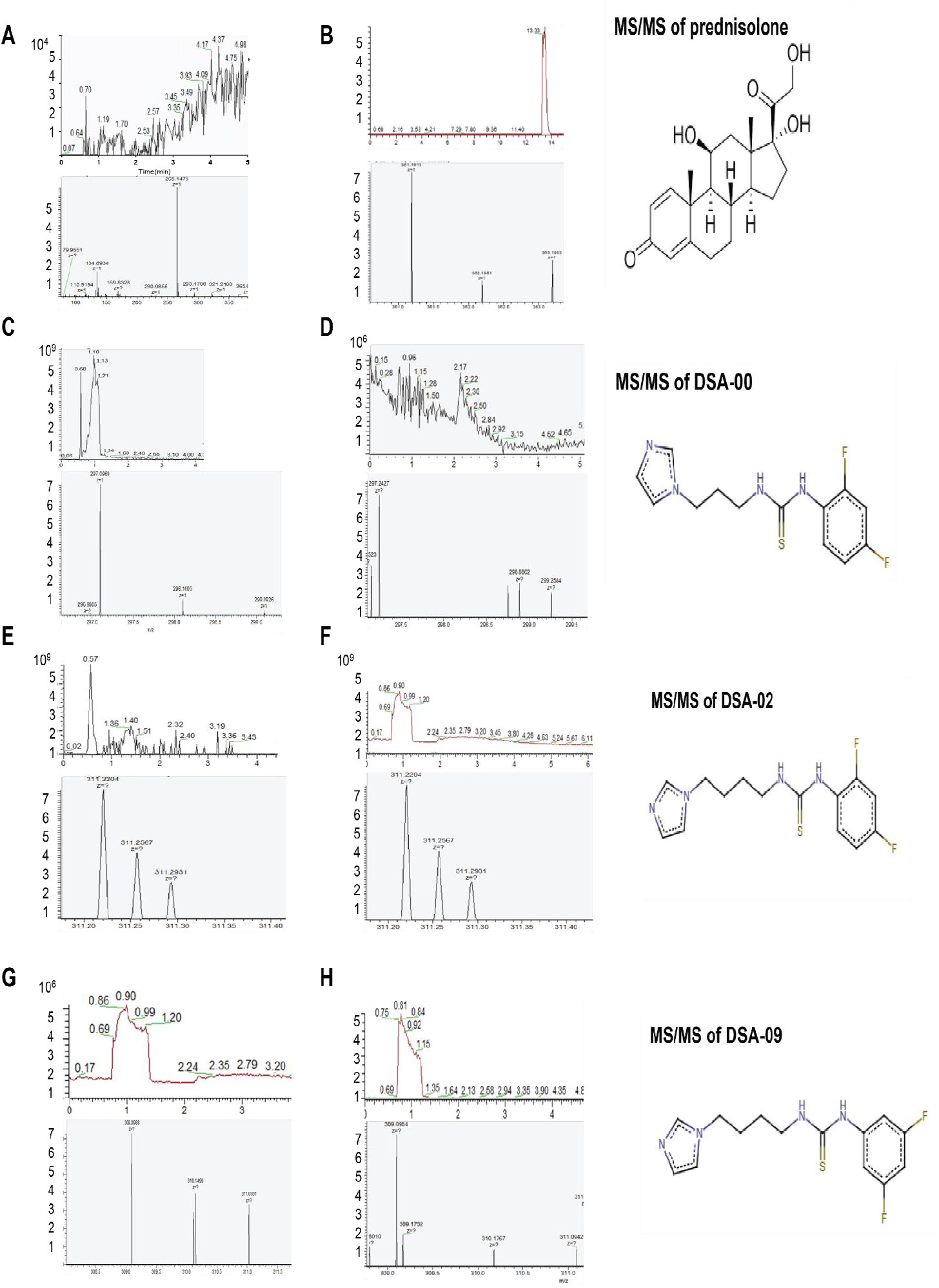
Typical MRM chromatograms of Thiourea derivatives and prednisolone (IS) in rat plasma. The blank sample(A), Blank sample with an internal standard (I.S.) prednisolone-spiked sample at 1000 ng/mL(B), DSA-00 spiked sample at 10000 pg/mL with an internal standard (I.S.) at 1000 ng/mL(C), plasma collected 2 hours after oral administration of aqueous DSA-00(D), DSA-02 spiked sample at 10000 pg/mL with an I.S. at 1000 ng/mL(E), plasma collected 2 hours after oral administration of aqueous DSA-02(F), DSA-09 spiked sample at 10000 pg/mL with an I.S. at 1000 ng/mL(G), and plasma collected 2 hours after oral administration of aqueous DSA-09(H).: The blank sample(A), Blank sample with an internal standard (I.S.) prednisolone-spiked sample at 1000 ng/mL(B), DSA-00 spiked sample at 10000 pg/mL with an internal standard (I.S.) at 1000 ng/mL(C), plasma collected 2 hours after oral administration of aqueous DSA-00(D), DSA-02 spiked sample at 10000 pg/mL with an I.S. at 1000 ng/mL(E), plasma collected 2 hours after oral administration of aqueous DSA-02(F), DSA-09 spiked sample at 10000 pg/mL with an I.S. at 1000 ng/mL(G), and plasma collected 2 hours after oral administration of aqueous DSA-09(H).

Further evaluating the recovery and matrix effect (Table S3) on the Quality Control (QC) levels Lower Limit QC (LLQC), Low QC (LQC), Medium QC (MQC), and High QC (HQC) was used to assess the analytical approach precision and accuracy. The findings, show that for both plasma and tissue homogenate samples, intra-day and inter-day precision, defined as Relative Standard Deviation (RSD), did not surpass 15%. Relative Error (RE), which measures accuracy, also came within the allowed range of 15% (Table S4). These results reveals that the analytical strategy utilized in this investigation for determining thiourea derivatives levels in rat plasma and tissue homogenates is repeatable and accurate and satisfies the preset acceptance criteria.

### 3.3. Pharmacokinetics of thiourea derivatives

After successful validation of method, examined the pharmacokinetics of thiourea derivatives in rats using orally and intravenous administration (10mg/kg). Then observed the mean plasma concentration-time profiles of DSA-00, DSA-02, and DSA-09 as well as calculated important pharmacokinetic parameters (Figure 3, Table 1, S5). The AUC_0-t_ (pg.h/mL) values for DSA-00, DSA-02, and DSA-09 groups were 6418.33 ± 156, 6428.44 ± 125.8, and 6769.25 ± 99.5, respectively. Therefore, oral bioavailability of thiourea derivatives viz DSA-00, DSA-02, and DSA-09 were determined to be 14.55, 14.24, and 15.34 respectively. Observed half-life (T_1/2_) of thiourea derivatives were 32.25, 3.38, 3.18 respectively, except DSA-00 had 10 times greater half-life as compared to the other derivatives of thiourea derivatives. Thiourea derivatives had mean residence time (MRT_0-∞)_ were 0.72, 0.69, 0.92, in DSA-00, DSA-02, and DSA-09 respectively. Thiourea derivatives clearance (CL) was lower in the DSA-02 (0.2), DSA-09 (0.2) than it was in the DSA-00 (0.5) (Table 1).

**Table 1:** Pharmacokinetic parameters of Thiourea derivatives in rat plasma.

| **Parameters** | **DSA-00** | **DSA-02** | **DSA-09** |
| --- | --- | --- | --- |
| **t<sub>1/2</sub> (h)** | 32.25 ± 0.5 | 3.38 ± 0.5 | 3.18 ± 0.22 |
| **CL (L/h/kg)** | 0.5 ± 0.05 | 0.2 ± 0.01 | 0.2 ± 0.01 |
| **MRT<sub>0-t</sub> (h)** | 0.80 ± 0.3 | 0.70 ± 0.2 | 0.93 ± 0.21 |
| **MRT<sub>0-∞</sub> (h)** | 0.72 ± 0.22 | 0.69 ± 0.14 | 0.92 ± 0.11 |
| **T max (h)** | 24 ± 5.25 | 24 ± 3.65 | 24 ± 9.56 |
| **C max (ng/mL)** | 4578.59 ± 56.36 | 4578.59 ± 56.36 | 4578.59 ± 56.36 |
| **AUC<sub>0-t</sub> (ng.h/mL)** | 6418.33 ± 156 | 6428.44 ± 125.8 | 6769.25 ± 99.5 |
| **AUC<sub>0-∞</sub> (ng.h/mL)** | 12477.50 ± 55.59 | 21211.61 ± 263.25 | 21661.59 ± 60.0 |
| **Bioavailability** | 14.55 ± .5 | 14.24 ± .5 | 15.34 ± .5 |

**Figure 3:**
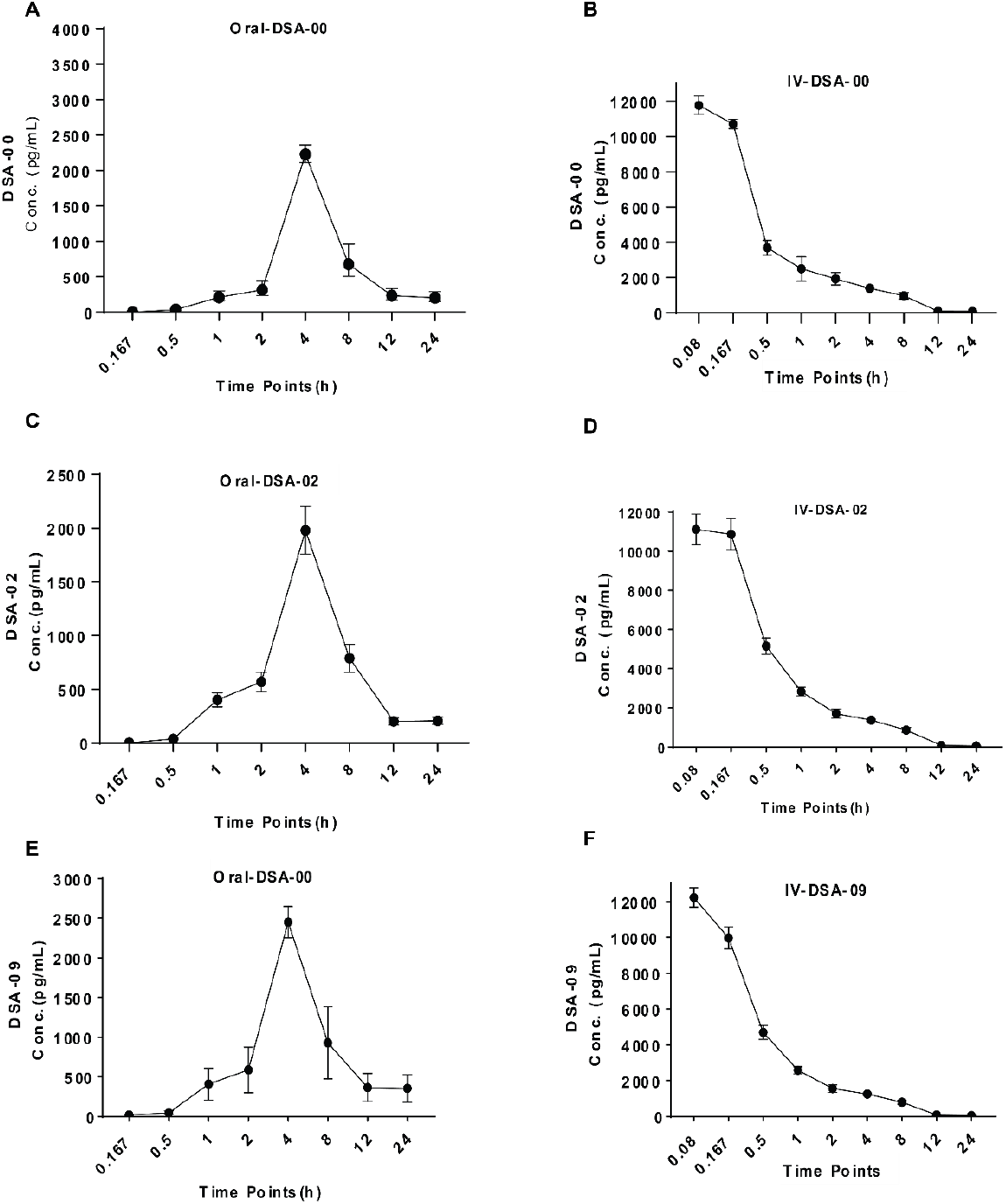
Plasma concentration of thiourea derivatives over a period of 24h in SD rat. Dose of 10mg/kg-bw of DSA-00 Orally and intravenously administration to SD rats, and collected blood from retro orbital route at different time points were collected and estimated the DSA-00 (A, B), DSA-00 (C, D), and DSA-00 (E, F) concentration in plasma on the basis of peak intensity via HPLC-MS/MS and analyzed by linear regression curve of concentrations vs peak intensity.

### 3.4. Tissue distribution of thiourea derivatives

Further method was used for the calculation of thiourea concentration in a variety of tissues, including heart, liver, spleen, lung, kidney, and brain homogenates. Following a single intragastric delivery of thiourea derivatives at a dose of 10 mg/kg, these tissues were processed at various time points (0.5, 1, 2, 4, 8, 12, and 24 hours). Thiourea derivatives tissue distribution patterns over time were examined (Figure 4), which showed that the substance was quickly absorbed and easily diffused across all of the examined tissues.

**Figure 4:**
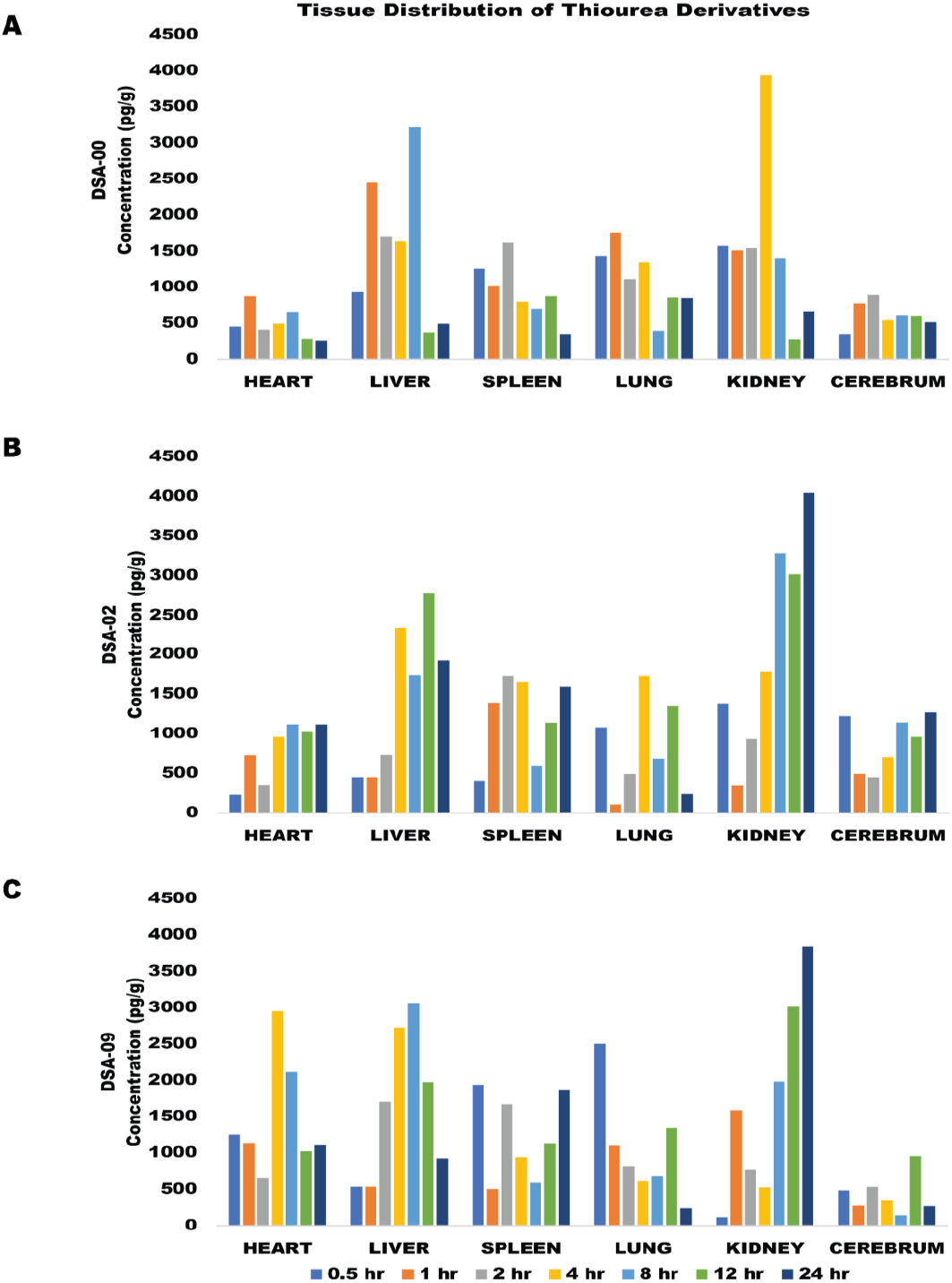
The concentration-time profile of thiourea derivatives viz. DSA-00(A), DSA-02(B), and DSA-09(C) in tissues following oral administration of 10 mg/kg-bw (n =6).

In all tissues, thiourea derivatives concentrations peaked at 1 hour after treatment and then significantly decreased over the course of the following 24 hours. This shows that none of the examined tissues significantly collect thiourea derivatives. The time to attain maximum concentration (T_max_) in the tissues was about 2-4 hours shorter in rats given a single intragastric dosage of thiourea derivatives (10 mg/kg), compared to plasma levels. The liver and Kidney have the highest maximum concentration of thiourea derivatives of all the tissues examined (Figure 4), followed by the heart, lung, spleen, and brain. This implies that thiourea derivatives might be especially effective at treating liver disorders and easily passed through the kidney. It’s noteworthy that thiourea derivatives was found in the brain following oral dosing, proving that it can cross the blood-brain barrier.

## 4. Discussion

Preclinical drug development requires drug pharmacokinetic and pharmacodynamic investigations via different delivery routes. Drug bioavailability and pharmacological properties depend on it. A sensitive, reproducible, and validated quantitative bio-analytical approach is essential for accurate, standardized pharmacokinetic analysis. Because of its sensitivity, LC-MS/MS is commonly used in pharmacokinetic studies. Moreover, recently some online tools available, like SwissADME online tool is a powerful research tool to predicts pharmacological properties of small molecules such as pharmacokinetics, bioavailability, drug-likeness, and medicinal chemistry friendliness (Daina et al., 2017). Here, we used swissADME tool, before performing in-vivo studies of thiourea derivatives. In order to support the computational prediction of the lipophilicity and polarity of the investigated small compounds, the graphical depiction of the Brain Or IntestinaL Estimated permeation technique (BOILED-Egg) has already been presented (Daina et al., 2017). As well, in a test trial conducted by us, thiourea derivatives were employed to prevent chronic Hepatitis B infection.(Singh et al., 2023; Kumar et al., 2023 communication) The prior study, which found that thiourea derivatives had similar effects to those of the well-known antiviral drugs (ETV), provides support for the current study assertion that thiourea derivatives may be more suitable than other non-nucleoside antivirals. Overall, the results suggested that thiourea derivatives would make good drug candidates because they resembled various ADME, bioavailability radar, and BOILED-Egg representations of well-known drugs (Figure 1).

SwissADME results revealed that thiourea derivatives have pharmacological properties. Therefore, these prediction results should be confirmed by an experimental, ADME, bioavailability, pharmacokinetics test and an in-vivo pharmacological assay for these lead thiourea derivatives. Thus, a robust and precise LC-MS/MS bio-analytical technique was developed and validated according to FDA and European Medicines Agency (FDA, 2021; EMEA, 2011). This approach quantifies thiourea derivatives in rat plasma and tissue homogenates. It’s offer sensitive, accurate, and specific for analysis. This innovative LC-MS/MS method improves drug pharmacokinetic evaluation over earlier methods. Furthermore, it was shown that thiourea derivatives matrix effect varied from 90.62% to 114.08% in tissue homogenate and plasma samples. These findings imply that the samples matrix did not significantly impede the analysis of thiourea derivatives. The e matrix effect of DSA-00, DSA-02, and DSA-09 in plasma samples ranged between 9.62 and 114.8 %, indicating that matrix did not interfere significantly with DSA-00, DSA-02, and DSA-09 analysis (Table S3).

Plasma concentrations of drugs aims to elicit the desired pharmacological response and reverse illness. Such pharmacological response is regulated by systemic circulation drug concentrations and drug availability at the receptor site its termed bioavailability of that drug (Chan and Holford 2001; Smolen 1978). In clinical practice, the time it takes to reach peak dosage (T_max_) is important because it varies from person to person and can be affected by its route of administration (e.g., orally, and intravenously). The half-life (t_1/2_) of a drug is often linked to T_max_. Short-lived drugs reach their peak quickly and are quickly eliminated, so they need to be taken more often to keep therapeutic amounts in the effective range. The highest concentration (C_max_) of a drug is linked to a key parameter called the area under the concentration-time curve (AUC). When a drug is eliminated in a linear way, the AUC is directly related to the amount of the drug that was taken. When drug clearance goes up, the AUC goes down. This means that the drug stays in the systemic circulation for less time, which makes the plasma drug levels drop more quickly. So, when clearance goes up, both the amount of drugs taken by the body as a whole and the area under the concentration-time curve go down. Therefore, we also studied the pharmacokinetic properties of DSA derivatives, which is helpful in calculating the right dosage which guarantees the desired effect of a candidate drug and its bioavailability (Lappin and Stevens 2008; Haider et al. 2021). While DSA-02 and DSA-09 were cleared at similar rates, DSA-00 took a longer time consistent with linear pharmacokinetics (Table 1). All thiourea derivatives reached the C_max_ values in 4 hrs. Further, DSAs compounds exhibited better excretion (clearance rate) but showed lower bioavailability. Note that oral ETV is rapidly absorbed from the gastrointestinal system, reaching maximal plasma concentrations (C_max_) within 1 hr, while C_max_ of thiourea derivatives were ∼4hrs. Therefore, to improve their bioavailability, DSA derivatives need to be conjugated with some agent that can alter the plasma membrane fluidity and promote passive transcellular drug penetration, modulate tight junctions to permit enhanced paracellular diffusion, and active efflux transporter modification (Gomez-Orellana 2005). Importantly, modified TDF is a pro-drug with 1.6-fold higher bioavailability than ETV. ETV is eliminated through unchanged substance by renal excretion (Yan et al. 2006; Park et al. 2017). It is not surprising that the liver and kidneys had the greatest levels of thiourea derivatives given their considerable vascularization and blood flow. Thiourea derivatives primary metabolism is probably primarily carried out by the liver, and its excretion is primarily carried out by the kidneys. This result indicates that more investigation into the potential of thiourea derivatives as an effective medication that can target the central nervous system is necessary.

Therefore, further investigation is required to establish the long-term effect of thiourea derivatives on renal functions. Nevertheless, the present study provides a foundation for further research into the pharmacological effects of thiourea derivatives with more preclinical studies and clinical trials.

## Supporting information

https://drive.google.com/drive/u/0/my-drive/ph

## Acknowledgements

This work was supported by grants to VK from the Department of Biotechnology (DBT), Government of India (Grant No. BT/PR30082/Med/29/1341/2018) and J.C. Bose National Fellowship (Grant No. SR/S2/JCB-80)/2012) of the Department of Science and Technology, Government of India, New Delhi JK PT and AKS received independent Senior Research Fellowships respectively from the Council of Scientific and Industrial Research, New Delhi and University Grants Commission, New Delhi for the period of this study.

## Figure Graphical Abstract

Sketch of Thiourea derivatives (DSA-00) followed evaluation of pharmacokinetics parameters by in-silico (1), and in-vivo, finally calculated of Bioavailability.

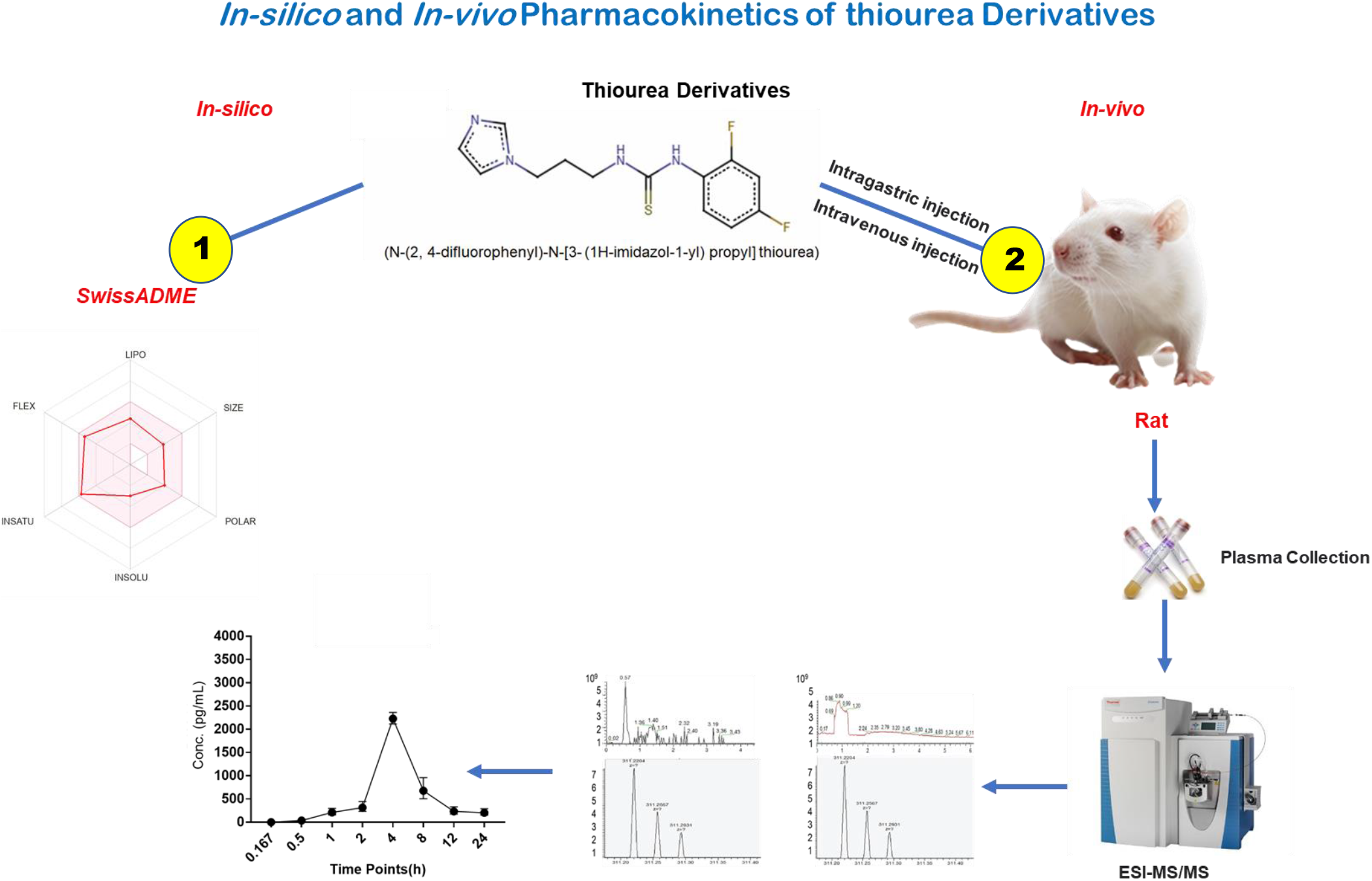

