## Supplementary material for "Analytical Method Development for Evaluation of Pharmacokinetics and Tissue Distribution of Thiourea-Based Antivirals through nLC/MS-MS": https://drive.google.com/drive/u/0/my-drive/ph

#### 1. Methods

1 Preparation of standard and quality control (QC) samples.

2 Chromatographic and mass spectrometric conditions

3 Experimental animal, collection of blood samples, and Pharmacokinetic by MS/MS

#### Tables

**Table S1**. Evaluation of pharmacokinetics, drug-likeness and medicinal chemistry friendliness of thiourea derivatives

**Table S2.** Summary of the LLOQ, linear ranges, regression equations, and association coefficients of analytes in rat plasma and tissue homogenate samples**.**

**Table S3.** Recovery and matrix effects on thiourea derivatives in plasma and tissues homogenate samples (n =6).

**Table S4.** The accuracy and precision of the method used to measure thiourea derivatives in biological materials (n=6).

**Table S5.** In vivo evaluation of pharmacokinetics parameters of thiourea derivatives

**Figures**

**Figure S1**. Chromatographic and MRM spectra of blank sample

**Figure S2.** Chromatographic and MRM spectra of blank spiked 100ng/mL sample

**Figure S3.** Chromatographic and MRM spectra of blank spiked 100ng/mL sample

**Figure S4.** Chromatographic and MRM negative mode of spectra of DSA-00 sample

**Figure S5.** Chromatographic and MRM positive mode of spectra of DSA-00 sample

**Figure S6.** Chromatographic and MRM positive mode of spectra of DSA-00 at the 10000ng/mL sample.

**Figure S7.** Chromatographic and MRM positive mode of spectra of DSA-00 at the 1ng/mL sample.

**Figure S8.** Chromatographic and MRM negative mode of spectra of DSA-02 sample.

**Figure S9.** Chromatographic and MRM negative mode of spectra of DSA-02 sample at 10000ng/mL.

**Figure S10**. Chromatographic and MRM negative mode of spectra of DSA-02 sample at 1ng/mL.

**Figure S11.** Chromatographic and MRM negative mode of spectra of DSA-09 sample.

**Figure S12.** Chromatographic and MRM negative mode of spectra of DSA-09 sample at 1000ng/mL.

**Figure S13.** Chromatographic and MRM negative mode of spectra of DSA-09 sample at 1ng/mL.

### Methods

#### 1 Preparation of standard and quality control (QC) samples

Accurately prepared the standard stock solution of 10 mg/mL DSAs required dissolving the standard in acetonitrile in the proper quantities. Additionally, a 1000 ng/mL solution of prednisolone as IS was made in acetonitrile. In order to produce calibration standards, 100 μL of blank rat plasma was spiked with the appropriate amount of the standard solutions, resulting in final DSAs concentrations of 0.01, 0.1, 1, 10, 100, 1000, and 10000 ng/mL. DSAs QC samples at three different concentration levels (1.00, 10.0, and 1000 ng/mL) were prepared in the same way.

#### 2 Chromatographic and mass spectrometric conditions

In the chromatographic separation, a C18 column was used, and the mobile phase consisted of a mixture of 30 volumes of solvent A (water) and 70 volumes of solvent B (methanol). The separation was performed using isocratic elution. The injection volume was 20 μL, and the flow rate was 0.6 mL/min. A divert valve was employed to direct the eluent to waste from 0 to 15 minutes and to the mass spectrometer (MS) from 10 to 15 minutes during each analysis run, which had a total duration of 15 minutes.

The mass spectrometer was operated in both negative and positive electrospray ionization (ESI) scan modes. The quantification of DSAs (thiourea derivatives) and the internal standard (IS) was achieved using multiple-reaction monitoring (MRM) transitions, specifically m/z 297.2, 311.2 and 309.09 for DSAs and m/z 361.19 for IS. The MRM settings were optimized for the collision gas ($N_2$) at 0.35 Pa, nebulizer at 30 psi, gas flow at 10 L/min, and gas temperature at 300°C. These methods adapted form (Sharma et al., 2022)

### 3 Experimental animal, collection of blood samples, and Pharmacokinetic by MS/MS

Approval was obtained from the Institutional Animal Ethical Committee (IAEC/ILBS/20/09) prior to the study. Six male rats weighing 300 ± 50 g and female rats weighing 200 ± 20 g were obtained from the Department of Comparative Medicine, Institute of Liver and Biliary Sciences, New Delhi, India.

The rats were housed in a controlled environment with a temperature of 22 ± 2 °C, relative humidity of 50 ± 10%, a 12-hour light-dark cycle, and provided with ad libitum access to food and water. Before the experiment, the rats were fasted for 12 h with free access to water. Collected blood from isoflurane euthanized rat via retro-orbital vein at time points of 0.833, 0.167, 0.5, 1, 2, 4, 8, 12.0, and 24.0 h after oral administration of thiourea derivatives at 10 mg/kg-bw. The collected blood samples were centrifuged at 5000 rpm for 10 minutes, and the resulting plasma samples were transferred to clean tubes and stored at -20°C till used. The quantification analysis of pharmacokinetic parameters was performed using Xcaliber software from Thermo Scientific. This analysis allowed for the examination of various pharmacokinetic parameters based on the collected plasma samples.

Table S1: Evaluation of pharmacokinetics, drug-likeness and medicinal chemistry friendliness of thiourea derivatives

| Property | **Parameters** | **DSA-00** | **DSA-02** | **DSA-09** |
|---|---|---|---|---|
| **Physio-chemical** | Formula | $C_{13}H_{14}F_2N_4S_1$ | $C_{14}H_{16}F_2N_4S_1$ | $C_{14}H_{16}F_2N_4S_1$ |
| | MW | 296.34 | 310.37 | 310.37 |
| | No. of H-bonds Acceptor /Donor | 3/2 | 3/2 | 3/2 |
| | Molar Refractory | 77.25 | 82.05 | 77.25 |
| **Lipophilicity** | Log P o/w(iLOGP) | 2.51 | 2.66 | 2.65 |
| | **XLOG3P** | 1.81 | 2.16 | 2.16 |
| | WLOGP | 3.19 | 3.58 | 3.58 |
| | MLOGP | 1.87 | 2.12 | 2.12 |
| | SILICOS-IT | 3.14 | 3.51 | 3.51 |
| **Water Solubility** | Log S(ESOL) | -2.76 | -2.98 | -2.98 |
| | ESOL Solubility (mg/ml) | 5.12E-01 | 3.22E-01 | 3.22E-01 |
| | ESOL Solubility (mol/l) | 1.73E-03 | 1.04E-03 | 1.04E-03 |
| | ESOL Class | Soluble | Soluble | Soluble |
| | Ali Log S | -2.98 | -3.35 | -3.35 |
| | Ali Solubility (mg/ml) | 3.08E-01 | 1.40E-01 | 1.40E-01 |
| | Ali Solubility (mol/l) | 1.04E-03 | 4.51E-04 | 4.51E-04 |
| | Ali Class | Soluble | Soluble | Soluble |
| | Silicos-IT LogSw | -4.95 | -5.34 | -5.34 |
| | Silicos-IT Solubility (mg/ml) | 3.36E-03 | 1.41E-03 | 1.41E-03 |
| | Silicos-IT Solubility (mol/l) | 1.13E-05 | 4.53E-06 | 4.53E-06 |
| **Pharmacokinetics** | GI Absorption | Yes | High | High |
| | BBB permeable | Yes | Yes | Yes |
| | P-gp Substrate | No | No | No |
| | CYP450 1A2 inhibitor | Yes | Yes | Yes |
| | CYP450 2C19 inhibitor | Yes | Yes | Yes |
| | CYP450 2C9 inhibitor | No | No | No |
| | CYP450 2D6 inhibitor | Yes | Yes | Yes |
| | CYP450 3A4 inhibitor | No | No | No |
| | Log Kp (Skin permeation) | -6.28cm/s | -6.66cm/s | -6.28cm/s |
| **Drug likeness** | Lipniski | Yes; o violation | Yes; o violation | Yes; o violation |
| | Ghose filter | Yes | Yes | Yes |
| | Viber | Yes | Yes | Yes |
| | Egan | Yes | Yes | Yes |
| | Muegge | Yes | Yes | Yes |

| | | | | |
|---|---|---|---|---|
| | Bioavailability Score | 0.55 | 0.55 | 0.55 |
| **Medicinal chemistry** | PAINS | O alert | O alert | O alert |
| | Brenk | Thiocarbonyl group | Thiocarbonyl group | Thiocarbonyl group |
| | Lead likeness | Yes | No; 1 violation | No; 1 violation |
| | Synthetic accessibility | 2.51 | 2.57 | 2.57 |

Table S2: Summary of the LLOQ, linear ranges, regression equations, and association coefficients of analytes in rat plasma and tissue homogenate samples.

| **Sample** | **Calibration Curve (Equation)** | **Coefficient ($R^2$)** | **Ranges(pg/mL)** |
|---|---|---|---|
| DSA-00 from Plasma | y = 0.017+3.3e-005*x | 0.997 | 10-10000 |
| DSA-02 from Plasma | y = 0.017+3.3e-005*x | 0.996 | 10-10000 |
| DSA-09 from Plasma | y = 0.017+3.3e-005*x | 0.996 | 10-10000 |
| DSA-00 from Tissue | y = 1.017*x - 461.7 | 0.997 | 1-5000 |
| DSA-02 from Tissue | y = 1.017*x - 561.7 | 0.998 | 1-5000 |
| DSA-09 from Tissue | y = 1.017*x - 361.7 | 0.998 | 1-5000 |

Table S3: Recovery and matrix effects on thiourea derivatives in plasma and tissues homogenate samples (n =6).

| Samples | DSA-00, DSA-02, DSA-09 Conc. (ng/mL) | Extraction recovery (mean±SD) of DSA-00, DSA-02, DSA-09 | Matric effect (mean±SD) of DSA-00, DSA-02, DSA-09 |
|---|---|---|---|
| Plasma | 10 | 74.55±11.85 | 101.68±9.77 |
| | 100 | 88.72±10.91 | 107.32±13.18 |
| | 2000 | 89.44±11.36 | 102.71±5.72 |
| Heart | 5 | 79.9±9.10 | 99.31±8.79 |
| | 200 | 79.7±4.40 | 100.88±3.09 |
| | 3000 | 78.5±10.70 | 100.38±14.30 |
| Liver | 5 | 96.89±6.19 | 100.57±12.41 |
| | 200 | 97.03±10.05 | 104.08±4.92 |
| | 3000 | 85.79±10.74 | 102.39±7.02 |
| Spleen | 5 | 82.33±8.40 | 100.56±12.11 |
| | 200 | 89.12±5.47 | 100.61±9.46 |
| | 3000 | 99.99±3.04 | 100.42±2.18 |
| Lung | 5 | 73.19±5.16 | 100.50±4.81 |
| | 200 | 76.52±6.81 | 99.92±12.51 |
| | 3000 | 72.41±12.10 | 98.31±8.43 |
| Kidney | 5 | 86.40±9.95 | 103.13±3.42 |
| | 200 | 81.03±6.55 | 109.98±3.54 |
| | 3000 | 96.45±5.00 | 100.82±1.66 |
| Cerebrum | 5 | 71.30±3.29 | 109.62±7.57 |
| | 200 | 74.26±1.79 | 103.93±9.74 |
| | 3000 | 92.60±9.80 | 106.85±13.33 |

Table S4: The accuracy and precision of the method used to measure thiourea derivatives in biological materials (n=6)

| Name | Spiked Conc. (ng/mL) | Measured conc. nLC-MS/MS (ng/mL) | Precision (RSD, %) | | Accuracy (RE, %) |
|---|---|---|---|---|---|
| | | | Intra-day (n=3) | Inter-day (n=3) | |
| DSA-00 | 1 | 0.99 ± 0.2 | 2.00 ± 0.5 | 2.30 ± 0.6 | 3.20 ± 0.9 |
| | 10 | 10.1 ± 0.3 | 4.45 ± 1.2 | 4.35 ± 2.4 | 5.09 ± 2.2 |
| | 1000 | 9987.01 ± 4.2 | 6.38 ± 2.2 | 6.38 ± 3.2 | 6.98 ± 3.4 |
| DSA-02 | 1 | 1.01 ± 0.3 | 0.9 ± 0.03 | 0.92 ± 0.6 | 1.20 ± 0.5 |
| | 10 | 9.97 ± 1.3 | 1.45 ± 1.2 | 2.35 ± 2.1 | 4.09 ± 1.2 |
| | 1000 | 9945.01 ± 15.2 | 6.78 ± 2.2 | 6.8 ± 3.2 | 5.98 ± 3.4 |
| DSA-09 | 1 | 1.1 ± 0.3 | 1.2 ± 0.02 | 1.92 ± 0.6 | 2.20 ± 0.5 |
| | 10 | 10.97 ± 1.3 | 3.45 ± 1.2 | 4.35 ± 2.1 | 6.09 ± 1.2 |
| | 1000 | 1012.01 ± 32.2 | 8.78 ± 2.2 | 9.8 ± 2.2 | 5.98 ± 3.4 |
| Plasma from DSA-00, DSA-02, and DSA-09 Treated | 1 | 1.07 ± 0.3 | 1.2 ± 0.02 | 1.92 ± 0.6 | 2.20 ± 0.5 |
| | 10 | 9.98 ± 1.3 | 3.45 ± 1.2 | 4.35 ± 2.1 | 6.09 ± 1.2 |
| | 1000 | 9997.5 ± 32.2 | 8.78 ± 2.2 | 9.8 ± 2.2 | 4.98 ± 3.4 |
| Tissue from DSA-00, DSA-02, and DSA-09 Treated | 1 | 0.8 ± 0.3 | 1.5 ± 0.2 | 2.92 ± 0.4 | 1.20 ± 0.5 |
| | 10 | 9.8 ± 1.3 | 2.45 ± 1.2 | 3.35 ± 1.1 | 4.09 ± 1.2 |
| | 1000 | 9987.5 ± 32.2 | 5.78 ± 3.6 | 10.8 ± 3.2 | 7.9 ± 3.4 |
| Prednisolone (I.S.) | 1000 | 1000.50 ± 0.52 | 1.50 ± 0.52 | 3.00 ± 5.1 | 1.60 ± 0.5 |

Table S5: In vivo evaluation of pharmacokinetics parameters of thiourea derivatives

| **S.No.** | **Pharmacokinetics Parameters** | **DSA-00** | **DSA-02** | **DSA-09** |
|---|---|---|---|---|
| **1.** | $C_{max}$ (pg/mL) | 16661.94 ± 15.25 | 13457.432 ± 25.36 | 19227.32 ± 178.69 |
| **2.** | $T_{max}$ (Minutes) | 24 ± 5.25 | 24 ± 3.65 | 24 ± 9.56 |
| **3.** | Intercept | 3.7 ± 0.5 | 3.7± 0.56 | 3.6 ± 0.5 |
| **4.** | Slope | 0.94 ± 0.003 | -0.11 ± 0.003 | 0.3 ± 0.001 |
| **5.** | Initial Conc. | 4578.59 ± 56.36 | 4578.59 ± 56.36 | 4578.59 ± 56.36 |
| **6.** | At zero time (C) | 4578.59 ± 56.36 | 4578.59 ± 56.36 | 4578.59 + 56.36 |
| **7.** | Dose (mg/Kg-bw) | 10.0 ± 0.05 | 10.0 ± 0.05 | 10.0 ± 0.05 |
| **8.** | Elimination K (hrs) | 0.22 ± 0.006 | 0.21 ± 0.005 | 0.21 ± 0.005 |
| **9.** | Distribution Vd (mL) | 218.41 ± 1.55 | 218.41 ± 1.55 | 218.41 ± 1.55 |
| **10.** | Distribution Vd (Ltrs) | 0.22 ± 0.01 | 0.22 ± 0.01 | 0.22 ± 0.01 |
| **11.** | Elimination T/1/2 (hrs) | 32.25 ± 0.5 | 3.38 ± 0.5 | 3.18 ± 0.22 |
| **12.** | Clearance (mL/hrs) | 47.7 ± 5.89 | 6.59 ± .9 | 7.2 ± 0.5 |
| **13.** | Clearance (Ltrs/hrs) | 0.5 ± 0.05 | 0.2 ± 0.01 | 0.2 ± 0.01 |
| **14.** | AUC(0-t) | 6418.33 ± 156 | 6428.44 ± 125.8 | 6769.25 ± 99.5 |
| **15.** | AUC(t-∞) | 12934.26 ± 55.59 | 21211.61 ± 263.25 | 21661.59 ± 60.0 |
| **16.** | Absolute Bioavailability (Oral) | 14.55 ± .5 | 14.24 ± .5 | 15.34 ± .5 |
| **17.** | Absolute Bioavailability (IV) | 2.41 ± .6 | 46.84 ± .6 | 2.43 ± .6 |

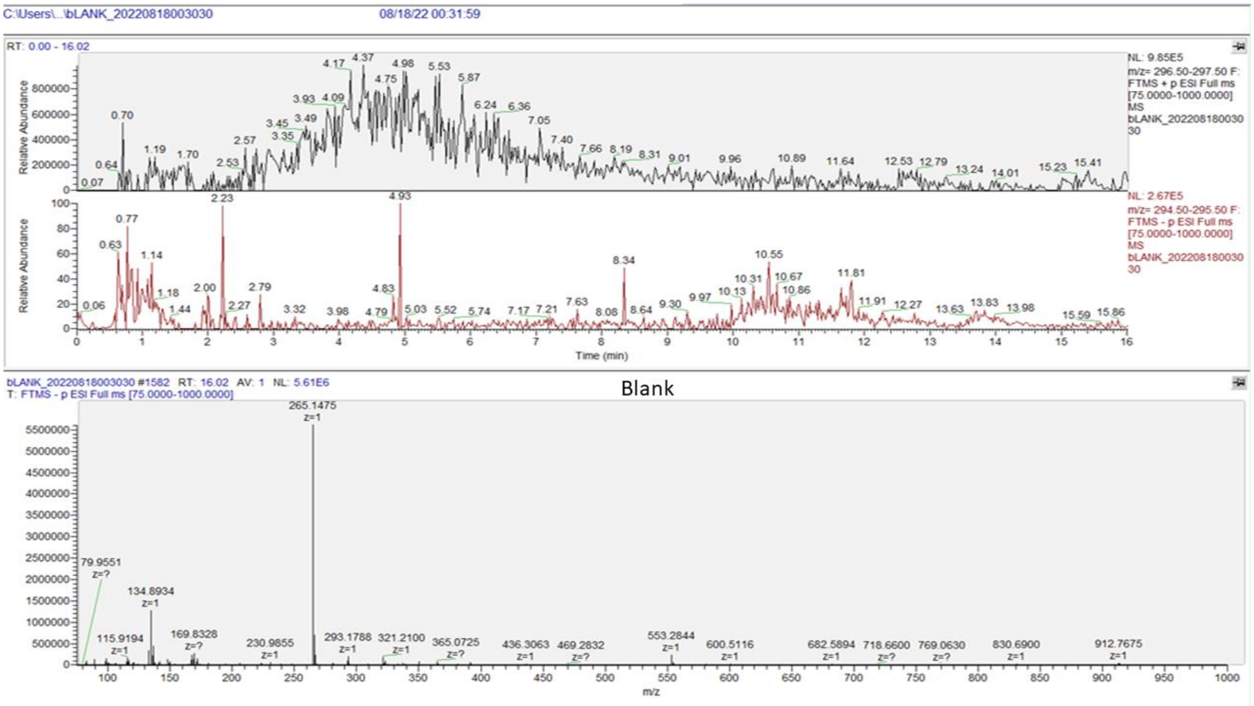

Figure S1. Chromatographic and MRM spectra of blank sample

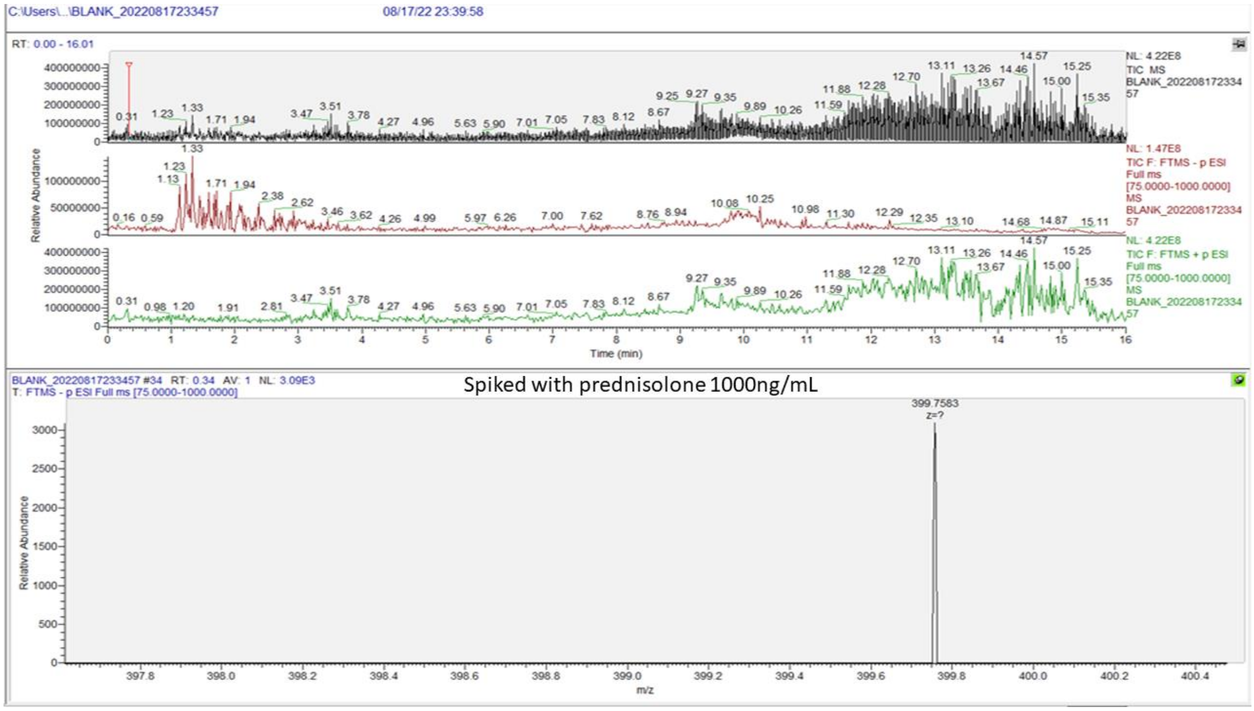

Figure S2. Chromatographic and MRM spectra of blank spiked 100ng/mL sample

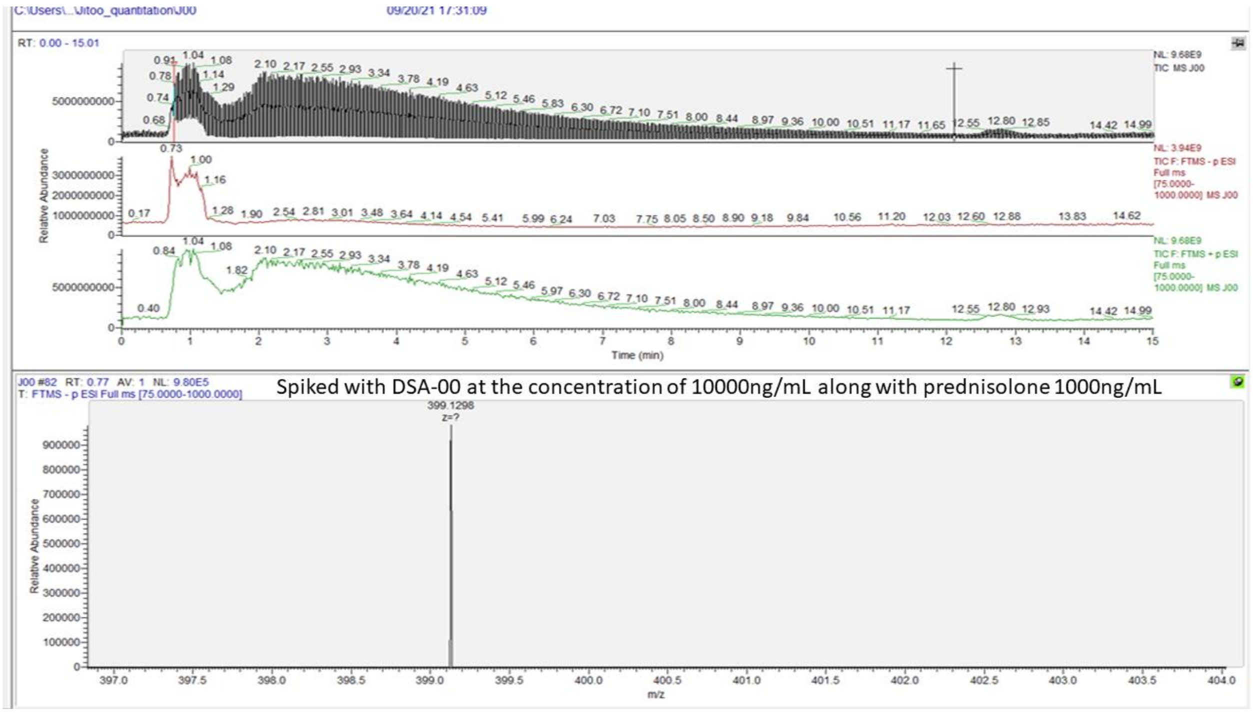

Figure S3. Chromatographic and MRM spectra of blank spiked 100ng/mL sample

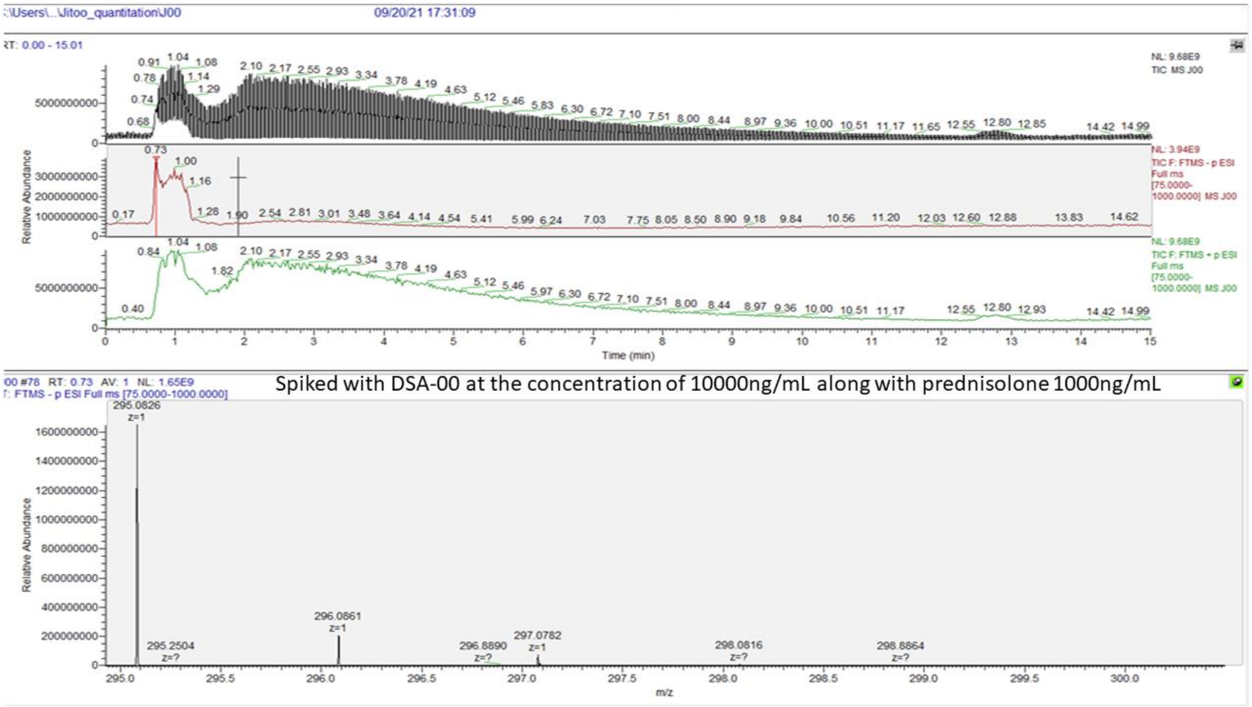

Figure S4. Chromatographic and MRM negative mode of spectra of DSA-00 sample

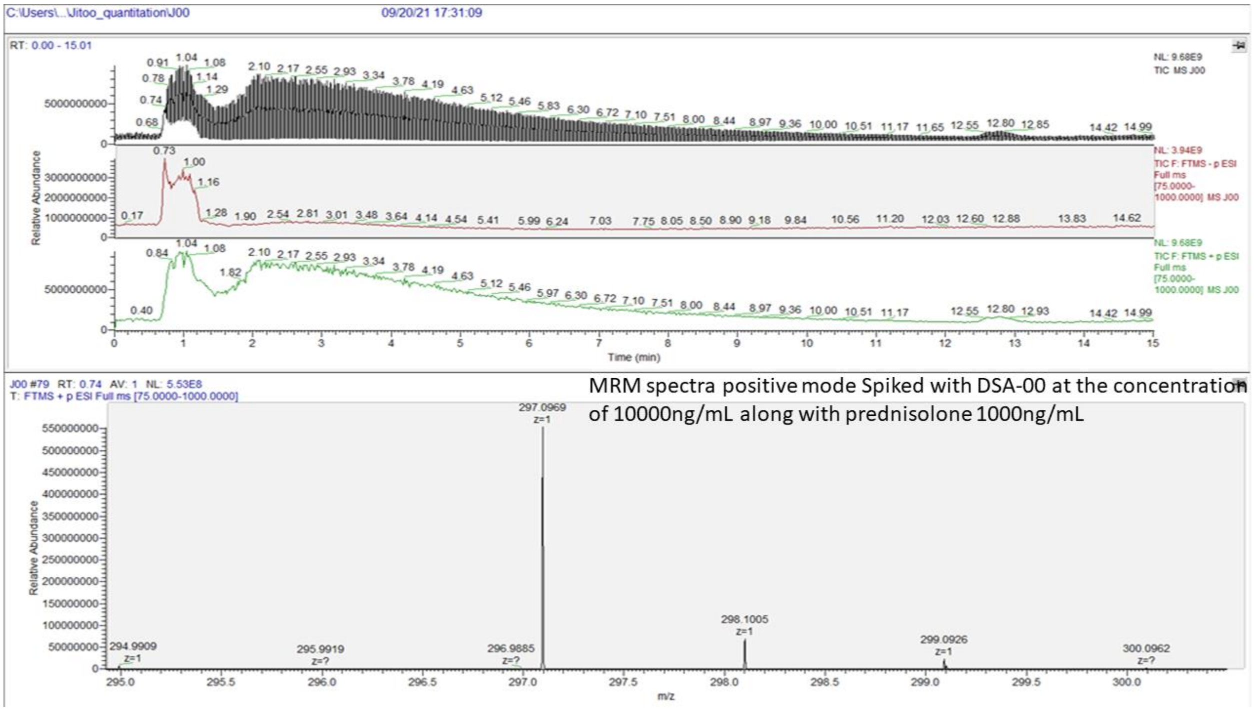

Figure S5. Chromatographic and MRM positive mode of spectra of DSA-00 sample

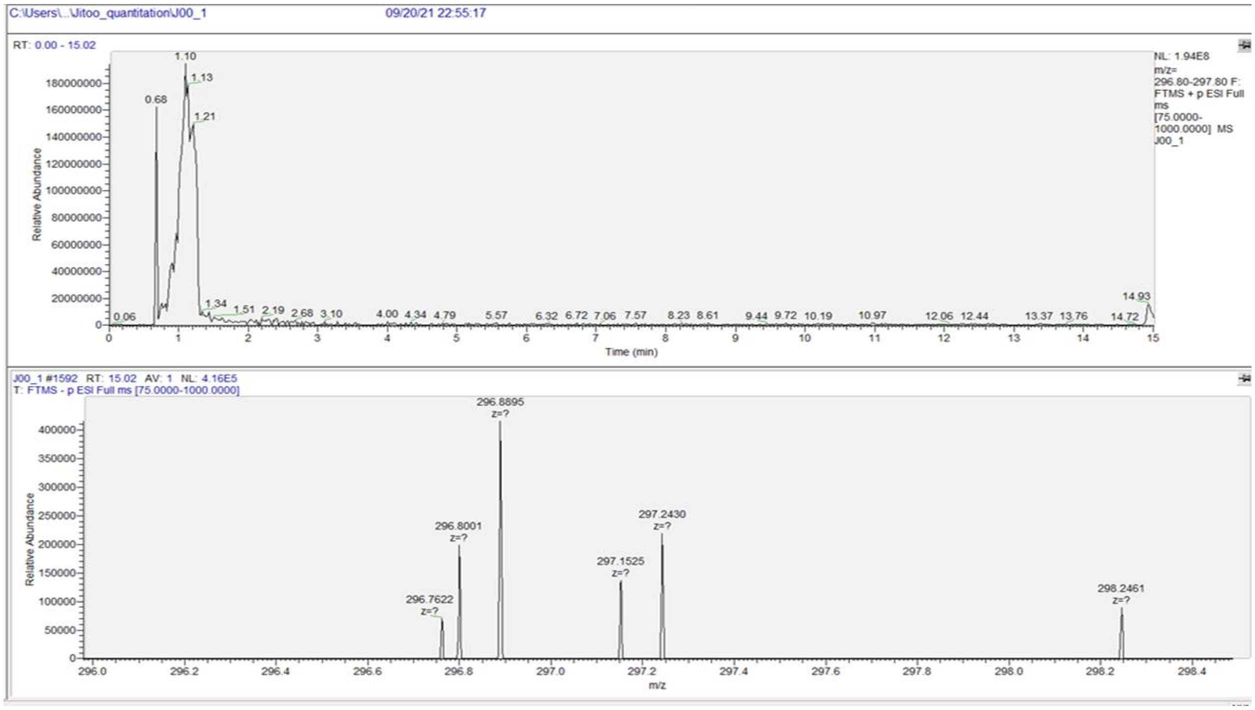

Figure S6. Chromatographic and MRM positive mode of spectra of DSA-00 at the 10000ng/mL sample.

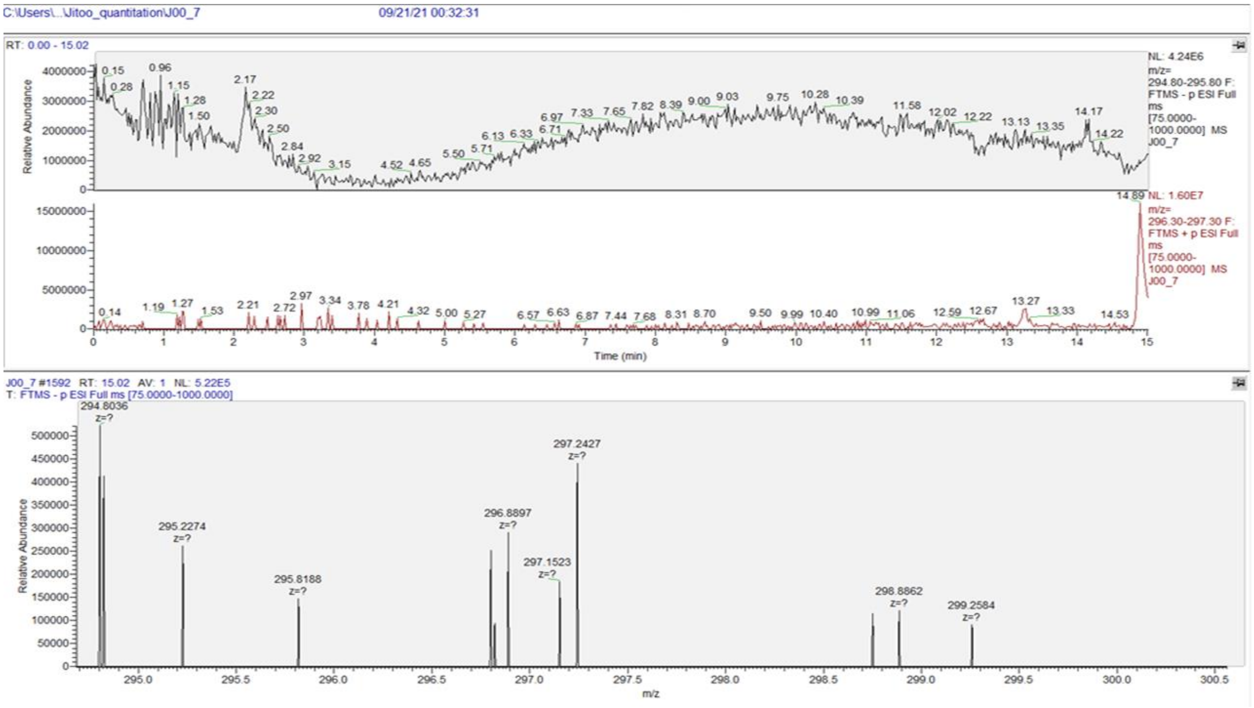

Figure S7. Chromatographic and MRM positive mode of spectra of DSA-00 at the 1ng/mL sample.

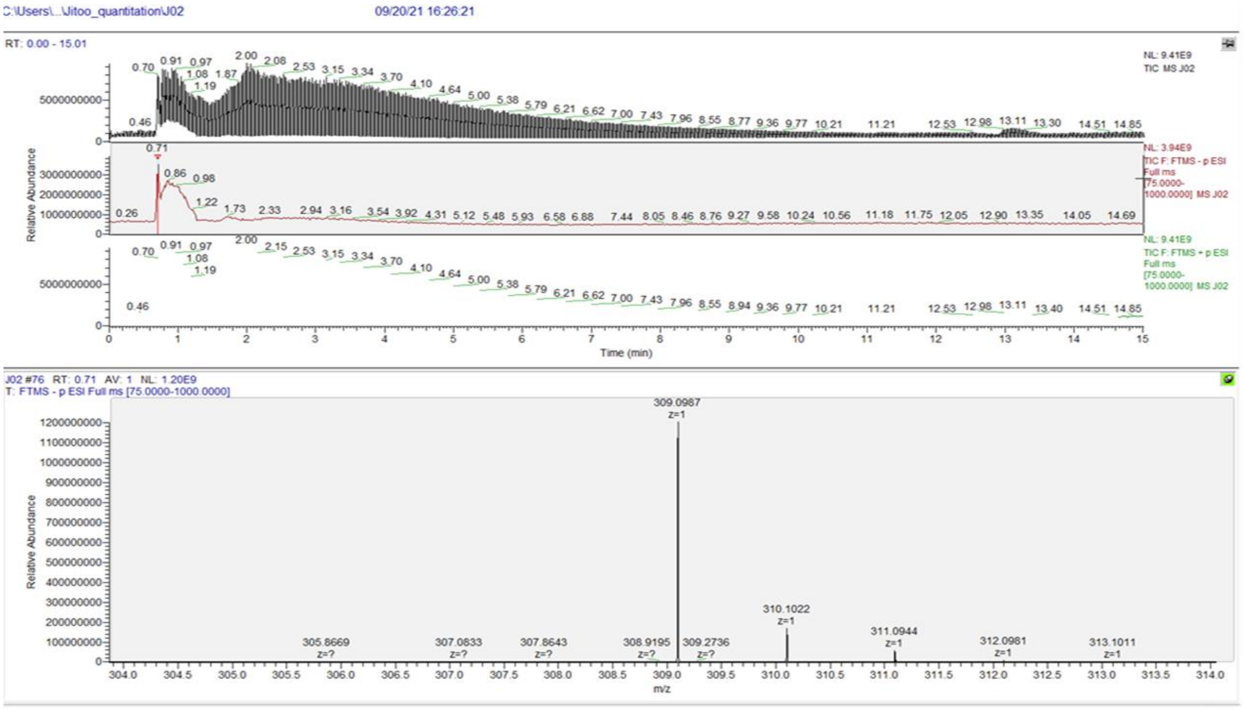

Figure S8. Chromatographic and MRM negative mode of spectra of DSA-02 sample.

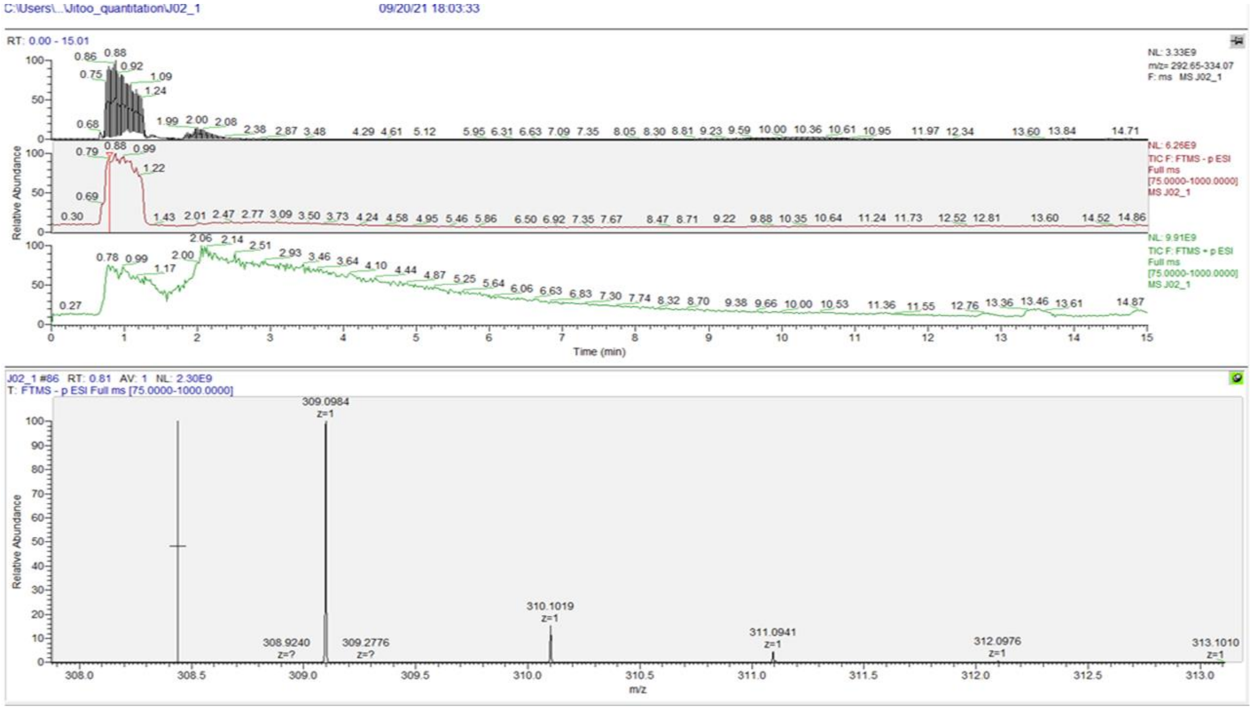

Figure S9. Chromatographic and MRM negative mode of spectra of DSA-02 sample at 10000ng/mL.

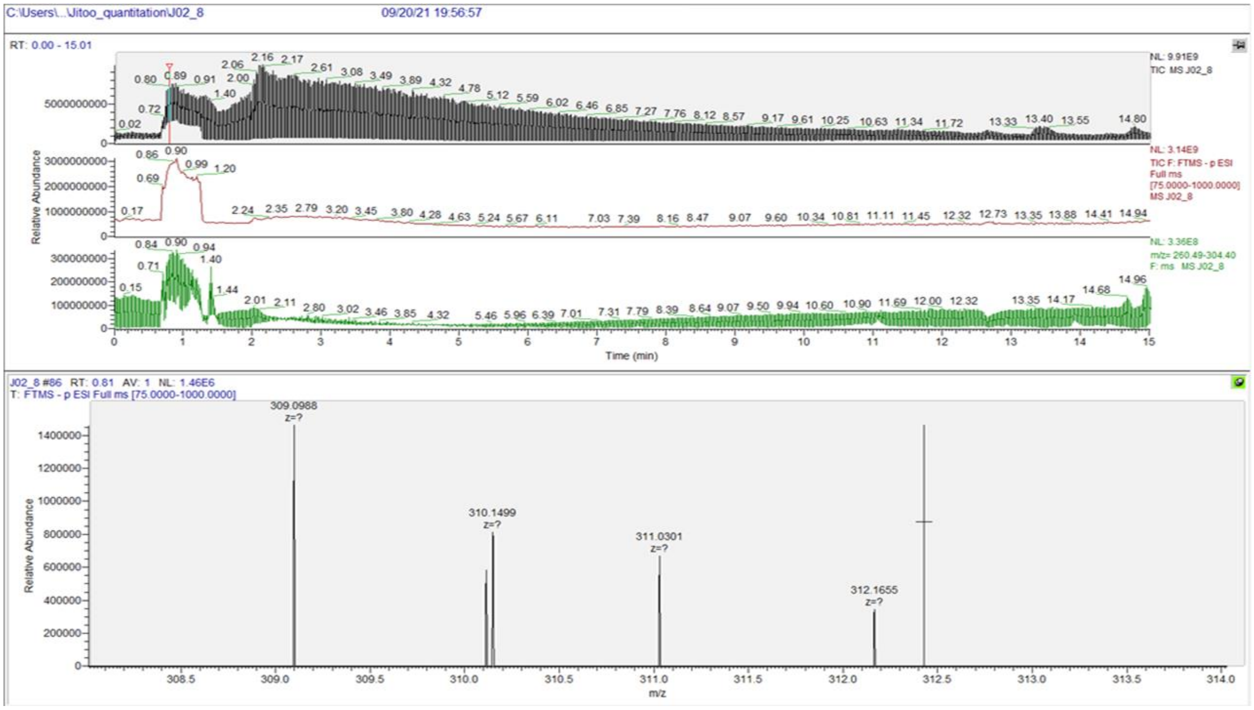

Figure S10. Chromatographic and MRM negative mode of spectra of DSA-02 sample at 1ng/mL.

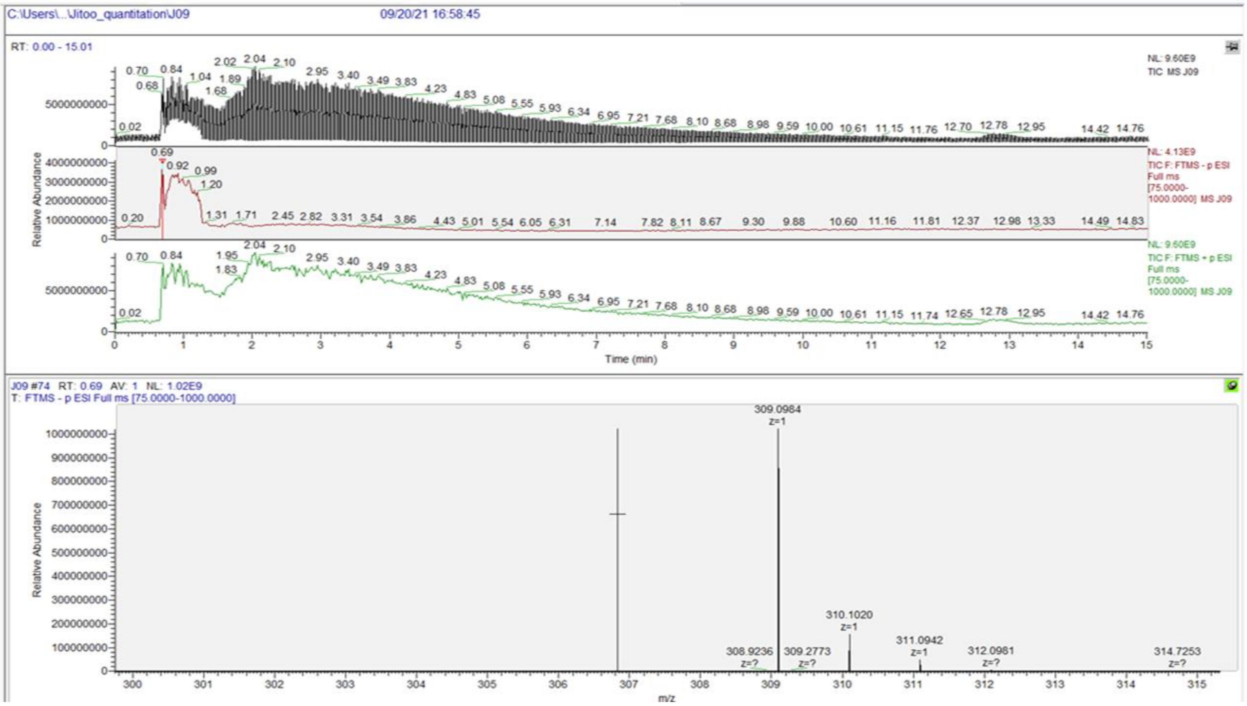

Figure S11. Chromatographic and MRM negative mode of spectra of DSA-09 sample.

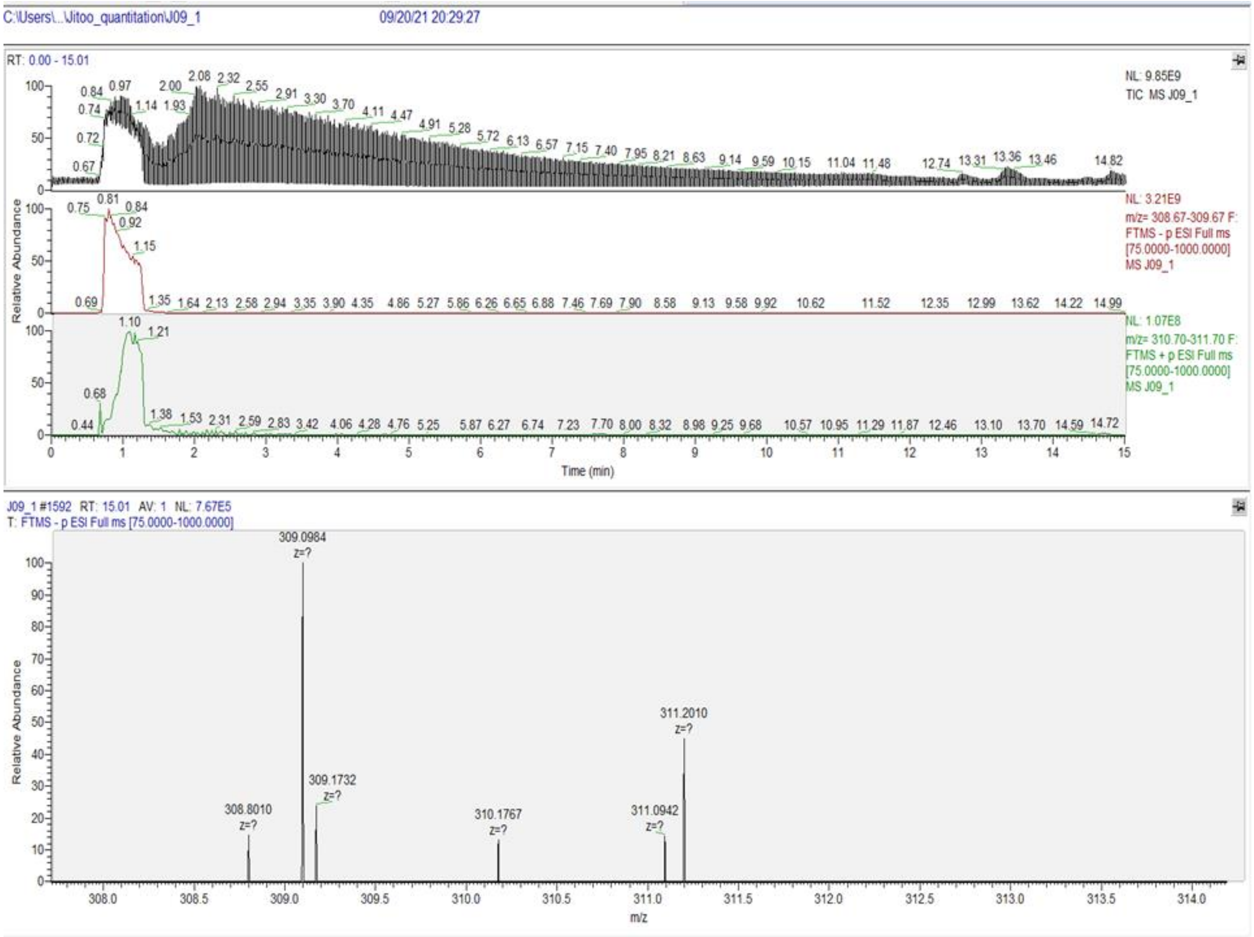

Figure S12. Chromatographic and MRM negative mode of spectra of DSA-09 sample at 1000ng/mL.

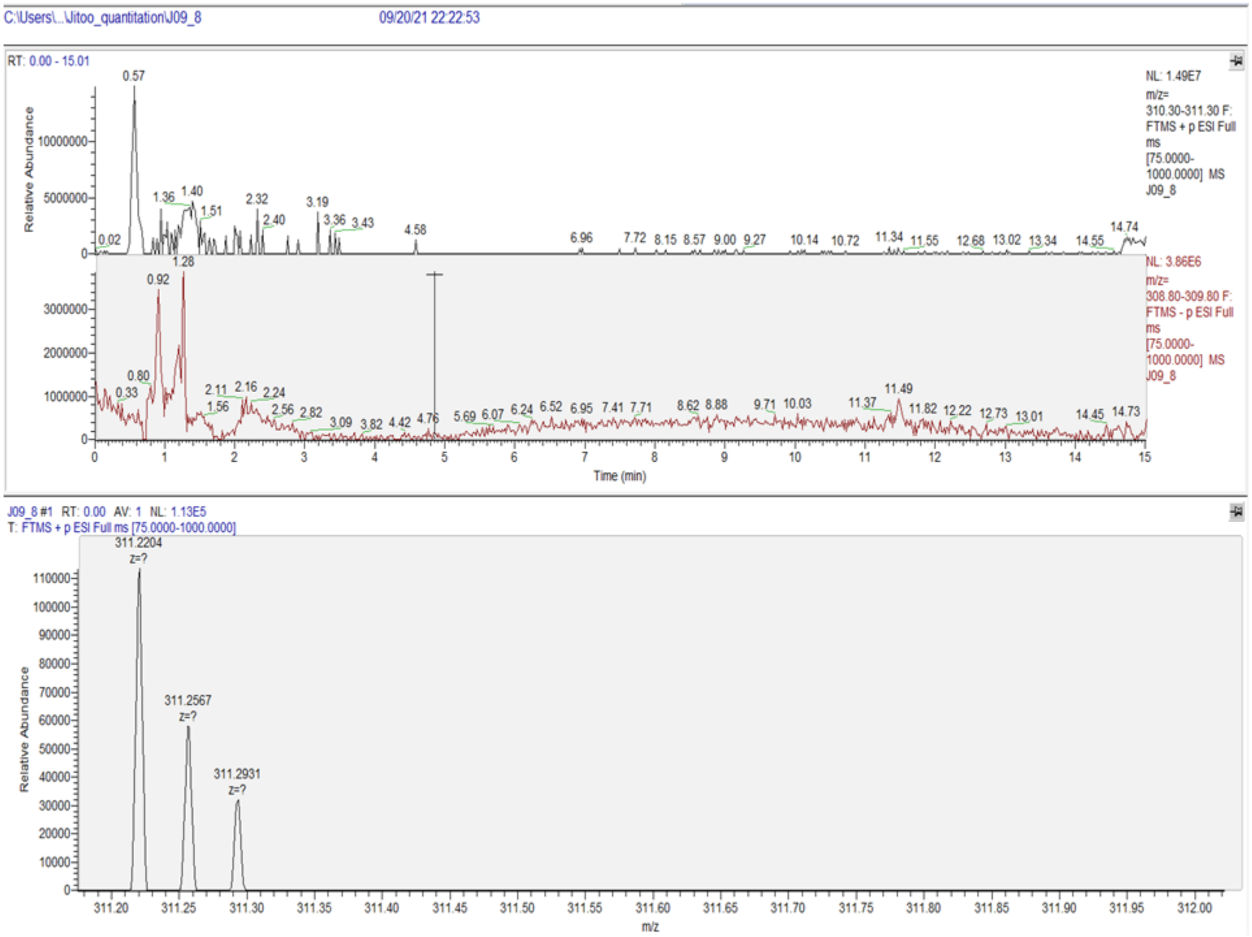

Figure S13. Chromatographic and MRM negative mode of spectra of DSA-09 sample at 1ng/mL.

Sharma, N., Bhat, S. H., Tripathi, G., Yadav, M., Mathew, B., Bindal, V., ... & Sarin, S. K. (2022). Global metabolome profiling of COVID-19 respiratory specimen using high-resolution mass spectrometry (HRMS). STAR protocols, 3(1), 101051.
